# Auxin fluctuation and PIN polarization in moss leaf cell reprogramming

**DOI:** 10.1101/2024.12.28.630436

**Authors:** Han Tang, Li-Hang Chen, Jiří Friml

**Affiliations:** Graduate Institute of Biochemistry, National Chung Hsing University, No. 145, Xingda Rd., South Dist., Taichung 40227, Taiwan, R.O.C; Institute of Science and Technology Austria (ISTA), Am Campus 1, 3400 Klosterneuburg, Austria; Advanced Plant and Food Crop Biotechnology Center, National Chung Hsing University, Taichung, 402, Taiwan

**Keywords:** Auxin, PpPINA, polarity, cell reprogramming, *P*. *patens*

## Abstract

Auxin and its PIN-FORMED (PIN) exporters are essential for tissue repair and regeneration in flowering plants. To gain insight into the evolution of this mechanism, we investigated their roles in leaves excised from *Physcomitrium patens*, a bryophyte known for its remarkable cell reprogramming capacity. We used various approaches to manipulate auxin levels, including exogenous application, pharmacological manipulations, and auxin biosynthesis mutants. We observed no significant effect on the rate of cell reprogramming. Rather, our analysis of auxin dynamics revealed a decrease in auxin levels upon excision, which was followed by a local increase before the reprogramming process began. Mutant analysis revealed that PpPINs are required for effective cell reprogramming, and endogenously expressed PpPINA-GFP accumulates polarly at sites that will develop into future filamentous stem cells. In addition, hyperpolarized PpPINA variants carrying mutated phosphorylation sites showed a marked delay in reprogramming, whereas endogenous or non-polar versions do not have this effect. These results underscore that both, the levels and the polarity of PpPINA are important for efficient cell reprogramming. Overall, these findings highlight the pivotal role of PIN polarity in plant regeneration. Furthermore, they suggest that understanding polarity mechanisms could have broader implications for improving regenerative processes across various plant species.

## Introduction

Regeneration is vital to repairing wounded tissues or replacing lost organs across animal and plant kingdoms. Only a few animals can regenerate a severed limb but the detached limb will never regenerate back to a new living body. In contrast, most plants possess relatively high plasticity to restore injured tissues and lost organs (Birnbaum and Alvarado 2008). Unlike animal regenerations, when a branch is detached from a plant, the detached branch usually regenerates and eventually re-develops into another individual plant. This unique characteristic of plants is not only fascinating from the mechanical point of view but also crucial for agriculture as many plants can be economically propagated by this method, which circumvents the necessity of seeding.

Different plant species show different regenerative capacities (Ikeuchi et al. 2016). Early-evolved, non-vascular bryophytes (mosses, liverworts, and hornworts) have higher regeneration capacity than vascular plants such as Arabidopsis (Sugimoto et al. 2019). In bryophytes, regeneration occurs in almost all wounded tissues, while in vascular plants, the regenerative capacity is limited in stem cell regions (Sugimoto et al. 2019). Differentiated somatic cells acquire the capacity to regenerate by cell reprogramming such that cells reset themselves to stem cells (Marhava et al. 2019). Some key components e.g. plant hormones, master transcription factors, and osmotic stress were identified to initiate this process (Sugimoto et al. 2019), but the molecular mechanisms are largely unknown.

Auxin, a versatile plant hormone that virtually contributes to every aspect of plant development and growth, is also crucial for plant regeneration. During most of the regeneration processes, auxin accumulates at the regeneration site to trigger the cell reprogramming (Hoermayer et al. 2020; Ye et al. 2020).

Despite the local biosynthesis of auxin, the auxin accumulation predominantly relies on its intercellular polar transport. In flowering plants, the auxin flow is mainly controlled via its PIN exporters (Adamowski and Friml 2015). The polarly localized PINs directly drive the direction auxin transport, which plays a dominant role in plant development ranging from embryogenesis to tropic directional growth of shoots and roots (Gallei et al. 2020).

Most studies in plant regeneration use the model flowering plant *Arabidopsis thaliana* as a working system. The regeneration of wounded vascular tissue is triggered by redistribution of PINs, which re-connect the auxin flow surrounding the cutting site (Hajný et al. 2022). The *pin1* mutant shows a lower regeneration rate (Sauer et al. 2006), suggesting an important role of auxin flow driven by polarized PINs in plant regeneration.

Little is known about the evolutionary conservation of the mechanisms underlying auxin signaling in plant regeneration. For this purpose, we study the early land plant moss, *Physcomitrium patens (P. patens)*. It has a short life cycle and grows mainly in filamentous cell types called protonema cells, along with tip growth and branches, from which few branch cells undergo cell fate transition and develop into leafy gametophores within 3 weeks (Rensing et al. 2020). The excised leaf from a gametophore is composed of a single-cell layer of leafy cells. These cells, when found next to the wounding site can reprogram into new filamentous stem cells. The regeneration takes place within 48 hours (Ishikawa et al. 2011). In *P. patens* several key components have been identified which play an important role in this process. STEMIN triggers cell reprogramming via chromatin modification (Ishikawa et al. 2019), CYCD1 is required for the cell cycle re-entry after cutting (Ishikawa et al. 2011), and PpCSP1, a homolog of a key transcription factor Lin28 for stem cell regeneration in animal cells, is indispensable for moss cell regeneration (Li et al. 2017). In addition to these critical factors, how and when the auxin and its exporter PINs are involved in this process is unclear.

In this study, we demonstrated that auxin has a moderate effect on moss leaf reprogramming and that the subcellular auxin level in a regenerative cell fluctuates after cutting displaying an initial decrease followed by an increase. We showed that *pin* knockout mutants exhibit significant defects in reprogramming, and PpPINA accumulates at the site of cell protrusion before cell expansion occurs. Further, the overexpression of hyperpolarized PpPINA variants significantly delayed reprogramming, whereas excess non-polar PIN proteins did not impact the reprogramming rate. These observations highlight the critical role of PpPINA’s proper polarity in cell reprogramming. Overall, our study provides insights into the roles of auxin and PIN proteins in plant cell reprogramming from an evolutionary perspective. Our results suggest that in addition to auxin export, PpPINA might serve as a crucial polarity marker, accumulating at the sites of future protrusions and sustaining the polarity during the outgrowth of reprogrammed stem cells.

## Results

### Changes in auxin level have a moderate effect on cell reprogramming

To investigate the role of auxin in moss leaf cell reprogramming, we first increased auxin levels by supplying various auxin forms throughout the entire reprogramming process. Detached leaves were placed on an agar medium enriched with different auxins, including Indole-3-acetic acid (IAA), naphthaleneacetic acid (NAA), and 2,4-dichlorophenoxyacetic acid (2,4-D). Despite these treatments, we observed no significant change in the reprogramming rate (Figure 1A).

**Figure 1.**
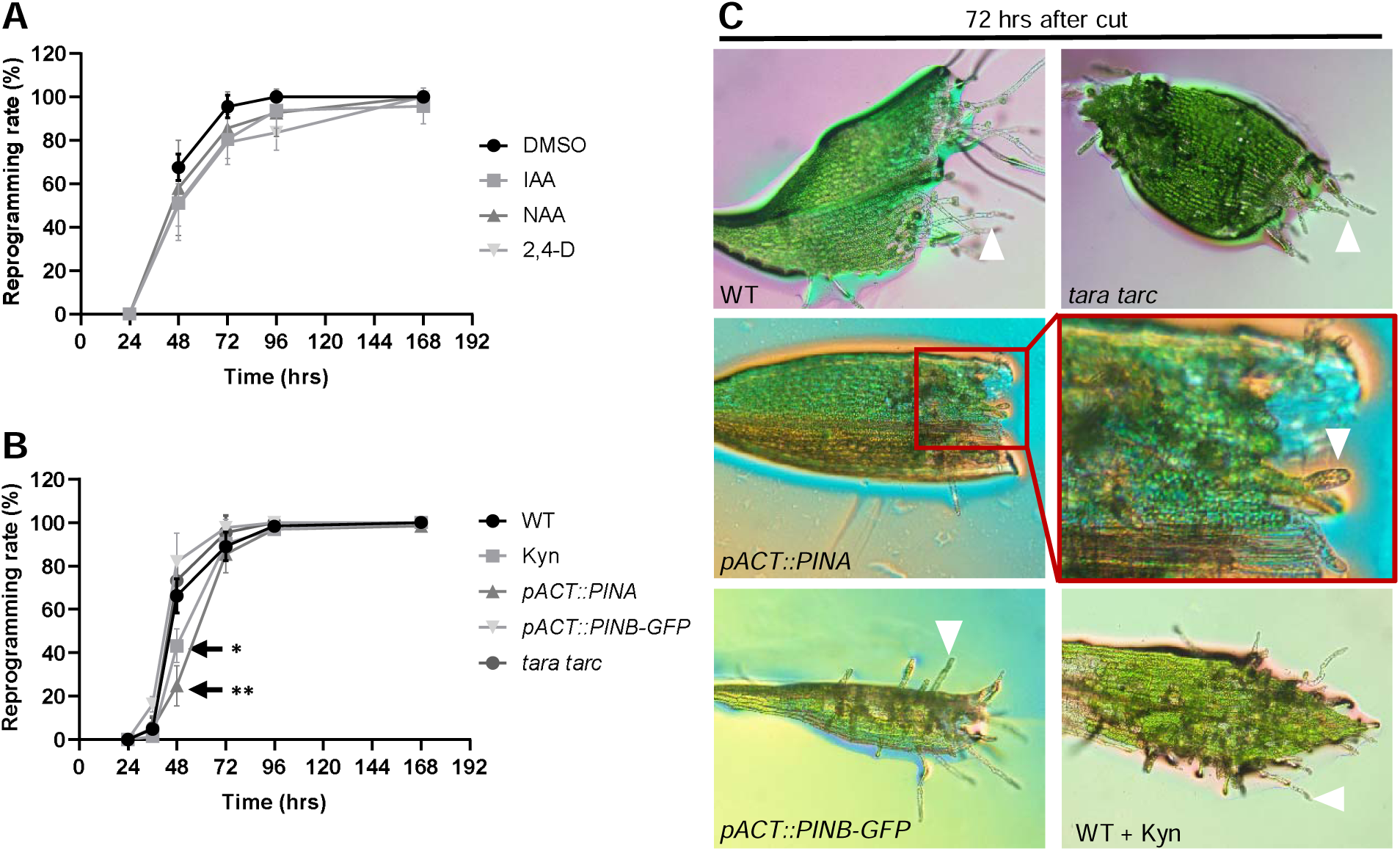
Manipulations of auxin level have a mild effect on moss cell reprogramming. (A) Reprogramming rates of detached leaves from wildtype (WT) treated with different analogs of auxin. The reprogramming rates were calculated by counting the percentages of detached leaves with at least one cell that appears to regenerate. N=3-4 independent experiment, n =20 leaves per experiment. The concentration of IAA, NAA, and 2,4-D is 10 μM for each. The same amount of DMSO was added to the growth medium as an internal control. (B) Reprogramming rates of detached leaves from WT with or without 10 μM L-kynuarine (labeled as kyn), and three indicated transgenic lines. Except for kyn treatment, all examined leaves were cultured on standard growth conditions. The delay of reprogramming at 48 hours in Kyn and pACT::PINA samples showed statistical significance compared to WT, students’ t-test. N=3-4 independent experiment, n =20 leaves per experiment. (C) Representative images of detached leaves from indicated lines. White arrowheads indicate protonema outgrowth, and the red frame presents delayed cell reprogramming in the pACT::PINA line.

Next, we explored the effects of reduced subcellular auxin levels using alternative approaches. We used transgenic plants carrying PpPINA or PpPINB-GFP driven by a constitutive promoter, both of which have been shown to present auxin depletion phenotypes in previous studies (Viaene et al. 2014). While overexpression of PpPINB-GFP revealed the same reprogramming capacity compared to wildtype (WT), PpPINA overexpression lines exhibited a one-day delay in cell reprogramming (Figure 1B, 1C). A similar delay phenotype was shown when treating excised leaves with L-Kynurenine, an inhibitor of auxin biosynthesis (Figure 1B, 1C). However, when we used the *tara tarc* double mutant, which lacks the key enzyme for auxin biosynthesis (Landberg et al. 2021), it showed no significant difference in reprogramming rates compared to the WT.

These findings suggest that excess auxin treatment has a relatively mild impact on moss cell reprogramming compared to other bryophytes and flowering plants. In *Marchantia polymorpha*, auxin supply significantly inhibits tissue regeneration (Ishida et al. 2022), whereas in flowering plants, varying ratios of auxin and cytokinin can accelerate tissue regeneration and root or shoot development (Ikeuchi et al. 2019; Xu 2018). Our results demonstrate a mild effect of auxin manipulation in moss leaf reprogramming, while the ratios of subcellular auxin and cytokinin in reprogramming cells were not simultaneously detected in this study. Our results indicate that *P. patens* may have developed an auxin-independent regulatory mechanism, in which cytokinin may play a role that contributes to its high reprogramming rate.

### Subcellular auxin levels fluctuate during cell reprogramming

Since changes in auxin levels did not affect reprogramming efficiency, we investigated whether subcellular auxin levels remained steady throughout the reprogramming process. To assess auxin at the cellular level, we utilized the *P. patens* R2D2 (PpR2D2) reporter line, which constitutively expresses mDII-nVENUS and DII-nTdTOMATO that respond rapidly to auxin in a dose-dependent manner (Figure S1). With the reporter constructs, elevated auxin levels lead to the rapid degradation of DII-nTdTOMATO, thus the ratio of mDII-nVENUS to DII-nTdTOMATO presents as an indicator of subcellular auxin levels (Thelander et al. 2019)

Initially, we tested the laser power for visualizing the R2D2 signal during reprogramming, while in our pretest, we found the signal was accumulated and highly saturated before reprogramming. To avoid the saturation that could interfere with the quantification, we chose to use lower laser power from the beginning of the time-lapse. We observed weak fluorescence signals from both MDII-nVENUS and DII-nTdTOMATO in excised leaves, which may be due to the weak laser power we exploited. However, 24 hours after cutting, the fluorescence signals became more distinct and visible (Figure 2A). This observation aligns with the activation of the key reprogramming regulator STEMIN which was shown to be activated after 24 hours and peaked at 30 hours (Ishikawa et al. 2019), indicating that the cell identity is transitioning from leaf to protonemata.

**Figure 2.**
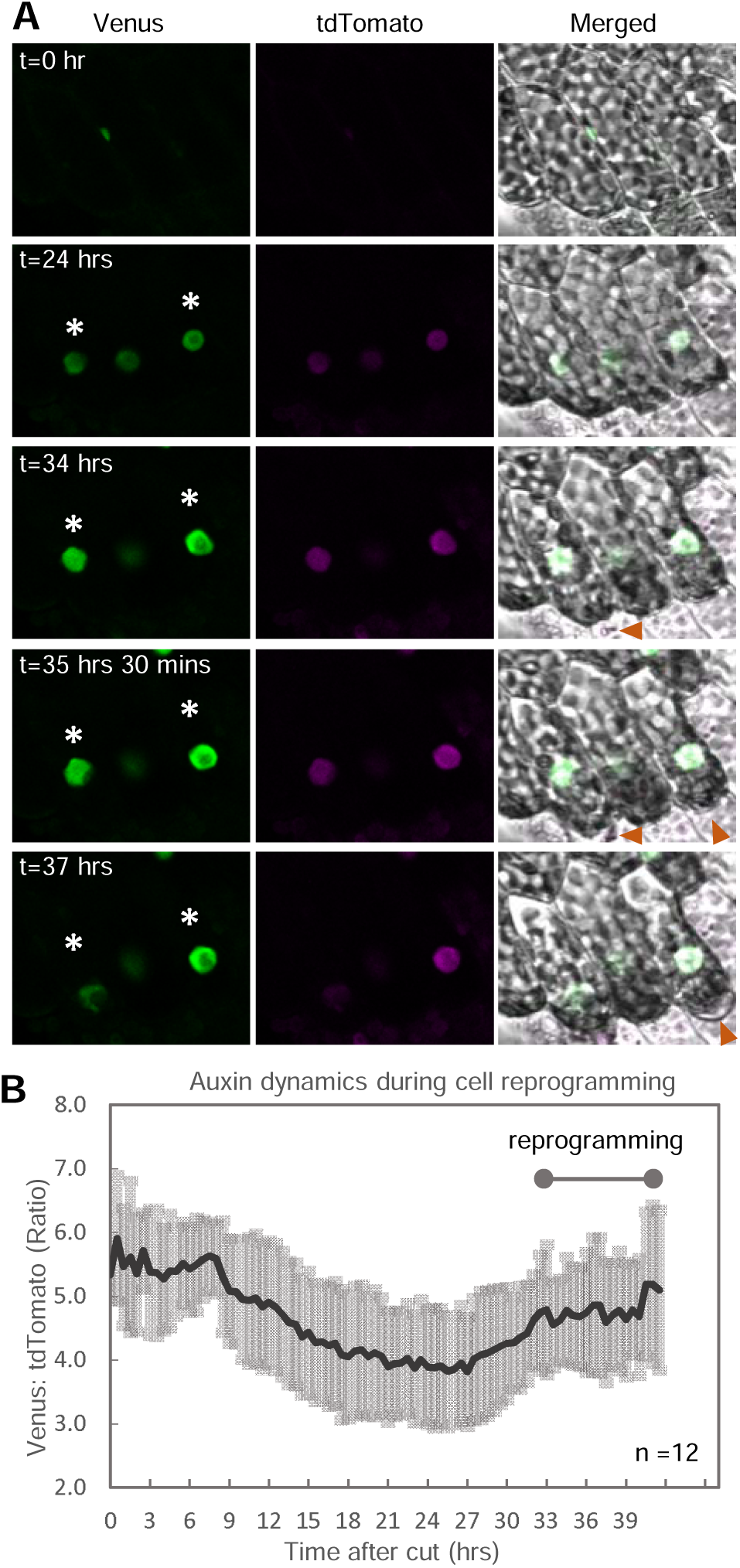
Subcellular auxin level fluctuates throughout moss leaf cell reprogramming. (A) Representative images of the auxin reporter R2D2 throughout cell reprogramming. The asterisks mark two reprogramming cells. Red arrowheads indicate the cell protrusion in the bright field. (B) The subcellular auxin level was measured and is presented based on the ratio of fluorescence Venus and tdTOMATO intensity. The intensity was tracked and time was measured from leaf excision to visible cell outgrow. The ratio represented auxin fluctuation throughout cell reprogramming.

We used the TrackMate function in Fiji to automatically track and quantify the fluorescence signals over time. Our data revealed that subcellular auxin levels began to decline approximately 9 hours after cutting, followed by a trend of increasing auxin levels around 24 hours after cutting, accompanying the reprogramming process (Figure 2B).

This detects, for the first time, the fluctuating subcellular auxin levels during cell reprogramming, however, it remains unclear whether this has a notable effect on reprogramming.

### PpPINA and PpPINB are required for cell reprogramming

In Arabidopsis, inflorescence stem regeneration following cutting relies on the redistribution of PIN proteins, which form auxin channels around the wounding site (Sauer et al. 2006). Mutants deficient in specific PIN proteins exhibit impaired vasculature regeneration, highlighting the critical role of PINs in this process (Mazur et al. 2020).

To explore whether PIN proteins are similarly essential for cell reprogramming in *P. patens*, we examined the reprogramming rates in *pina* and *pinb* single-gene knockout mutants, as well as in a *pina pinb* double knockout mutant. We first validated the auxin export capacity of examined *pin* mutants by isotope-labeled IAA. Fresh moss tissues were cultivated in a liquid medium supplied with H^3^-IAA for 24 hours, followed by washout and cultivation in a fresh medium for another 24 hours. The exported H^3^-IAA in the medium was detected by an isotope detector as described previously (Lewis and Muday 2009). Despite the *pina pinb* mutant exhibiting similar output as the single-gene knockout mutants, the auxin export assay showed that all examined *pin* mutants have a lower capacity of exporting auxin compared to WT (Figure 3A).

**Figure 3.**
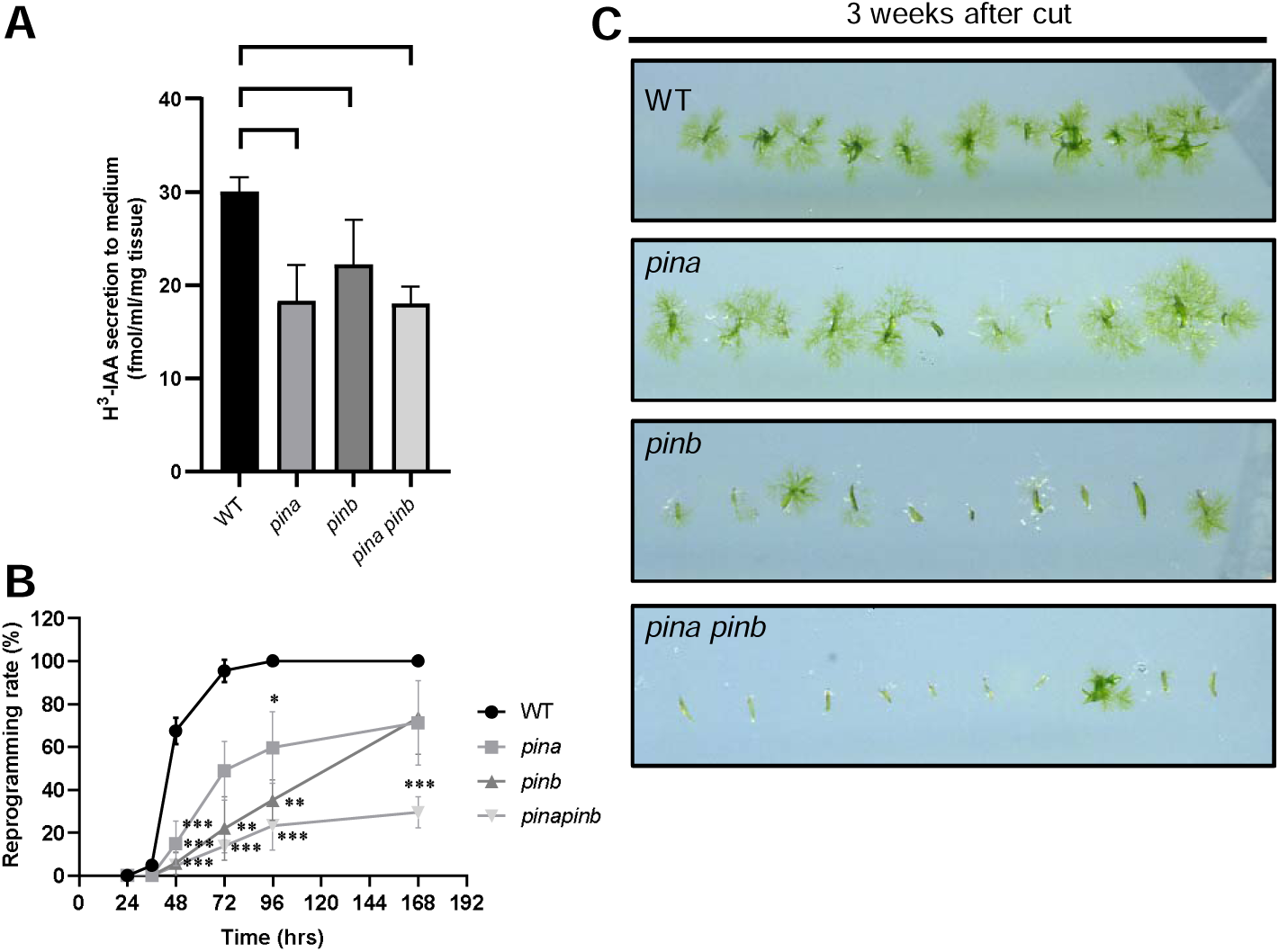
Auxin efflux carrier PIN proteins are required for moss leaf cell reprogramming. (A) Auxin export assay with wildtype and indicated *pin* mutants. Fresh tissues were cultivated on a liquid medium with H^3^-IAA for 24 hours followed by washout and cultivation in a fresh medium for an additional 24 hours. The amount of H^3^-IAA exported to the medium was detected as described (see methods). The *pin* mutants possess a lower capacity of exporting auxin compared to wildtype. Data were obtained from 6 independent experiments for each line. (B) Detached leaves from wildtype (WT) and indicated *pin* mutants were cultivated on the standard growth medium after three weeks of cutting. (C) Reprogramming rates of leaves detached from WT and indicated *pin* mutants. Single mutants *pina* and *pinb* showed delayed cell reprogramming, and the double mutant *pina pinb* exhibited the most severe defect in cell reprogramming.

We used the same mutants to examine their ability to regenerate from excised leaves. Our analysis revealed that single and double-knockout mutants had significantly reduced reprogramming capacity compared to WT (Figure 3B). Notably, the reprogramming process was not merely delayed but completely abolished in the double mutants. After three weeks, the leaves of these mutants remained viable but failed to reprogram into protonema (Figure 3C). Given that PIN proteins are functioning for auxin export, one might expect their knockout to lead to excessive auxin accumulation. The *pin* mutants used in this study have been shown to facilitate the cell fate switch in protonema development, which is opposite to the phenotypes while overexpressing PpPINA (Viaene et al. 2014). In line with this, the results of our assays with *pin* mutants showed a lower auxin export capacity (Figure 3A) and PpPINA-GFP overexpression showed a higher amount of auxin exported (Tang et al. 2024). The results showed that auxin export levels match the genotypes, supporting that *pin* mutants accumulate excessive auxin. Despite this, exogenous auxin supply did not impair cell reprogramming as the *pin* mutants did (Figure 1B, 3B). This suggests that PIN proteins may have additional roles in cell reprogramming beyond a simple auxin export.

### PpPINA polarization predicts the future site of cell outgrow

To explore the role of PpPIN proteins in cell reprogramming further, we observed endogenously expressed PpPINA fused with a green fluorescent protein (*proPINA::PpPINA-GFP*). Because our preliminary test showed that the signal of PpPINA-GFP was not visible within 24 hours after cutting, we started the time-lapse imaging from 30 hours after cutting. Multiple time-lapse movies were synchronized by the timing of cell outgrow, thus the time scale presented in Figure 4B was time before outgrow. Time-lapse imaging of PpPINA-GFP revealed its accumulation at the future protrusion site prior to the visible cell outgrowth (Figure 4A, 4B). Specifically, PpPINA-GFP began to polarize on the plasma membrane approximately 8 hours prior to cell protrusion, whereas non-regenerative cells displayed a uniform and low-intensity distribution of PpPINA-GFP (Figure 4B).

**Figure 4.**
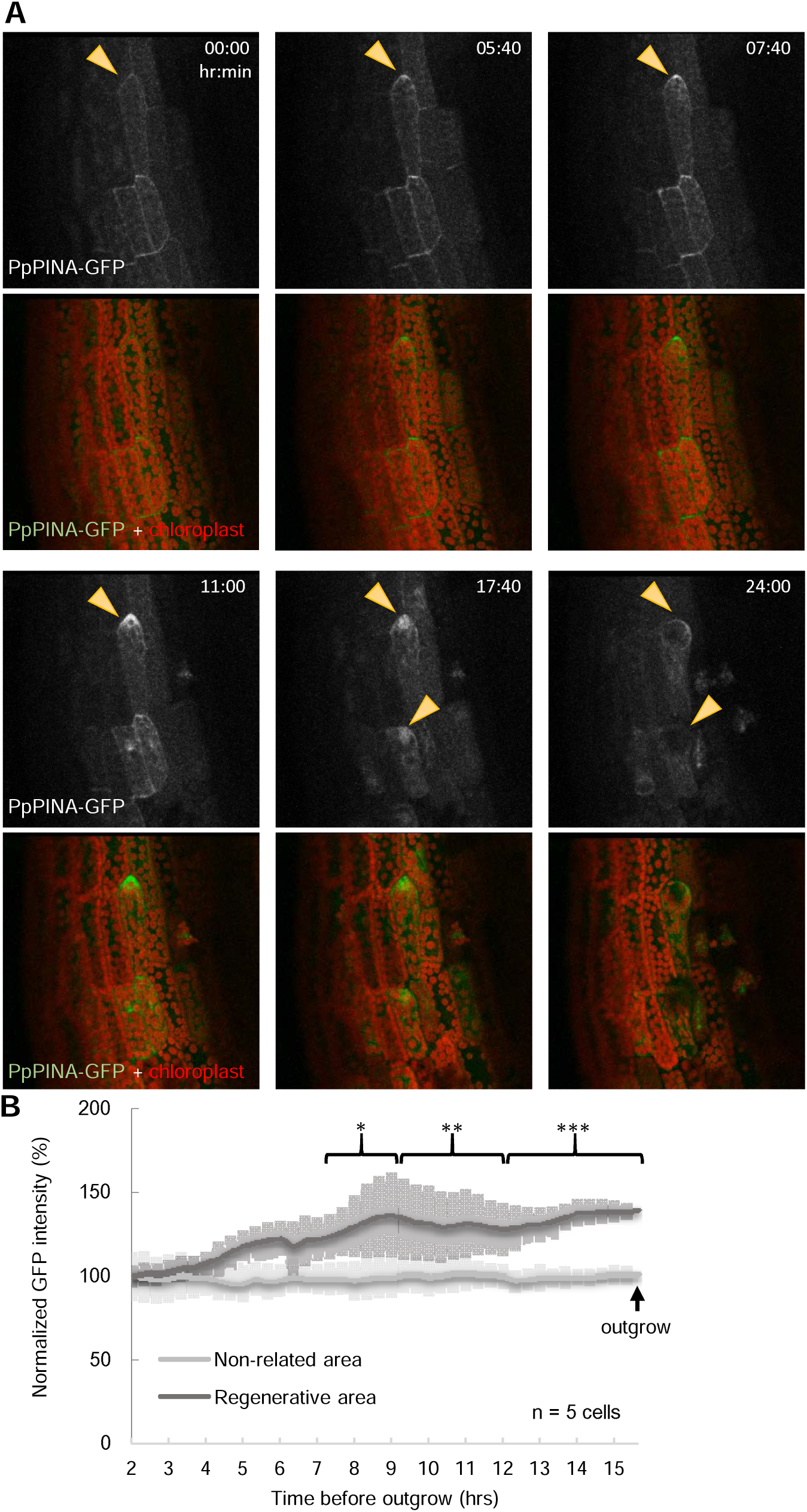
PpPINA is polarized at the protrusion site prior to cell outgrows. (A) Representative images of PpPINA-GFP polarization prior to visible cell outgrowing. Yellow arrowheads indicate the accumulation of the PpPINA-GFP signal. The signal diminished after the cell outgrew due to the elongation of reprogrammed cells that grew out of focus. Time zero indicated 30 hours after cutting, the time-lapse imaging was carried out from 30 to 54 hours after cutting. (B) Quantitative analysis of PpPINA-GFP intensity in the regenerative area near the reprogramming site. The average intensity was calculated. The time-lapse images were synchronized by the moment of cell outgrow. The intensity was normalized to the original intensity at the starting time point of imaging. As a control, a non-related area of comparable size distant from the regenerative sites was chosen.

It is noteworthy that, despite the more severe defects in cell reprogramming observed in the *pina pinb* double mutant, PpPINB was not detectable during the reprogramming process. This suggests a potential functional redundancy between PpPINA and PpPINB, with PpPINA playing a dominant role. Alternatively, PpPINB’s function may not be directly involved in regeneration. The observed polarization of PpPINA-GFP at the future reprogramming site prior to cell outgrowth led us to hypothesize that PpPINA may act as a polarity landmark for cell reprogramming.

### Overexpression of hyperpolarized PpPINA impedes cell reprogramming

To investigate whether PpPINA functions as a polarity landmark in cell reprogramming, we manipulated its polarity by overexpressing variant PpPINA with altered polarity in protonema cells (Figure 5A). An initial attempt to search for potential phosphorylation sites on PpPINA revealed several residues that are essential for the correct polar localization of PpPINA, as shown below. In WT background, we generated transgenic plants inducible overexpressing the PpPINA variants with point mutations at either single serine (PpPINA-GFP^S527A^) or two phosphorylation sites (T192A/S527A, referred to as PpPINA-GFP^AA^). Mutations on potential phosphorylation sites resulted in hyperpolarization of these PpPINA variants (Figure 5A), while WT PpPINA under the same inducible conditions was not hyperpolarized, indicating the hyperpolarization of PpPINA variants was attributed to their mutations rather than expression levels. To verify their auxin export ability, we performed an isotope-labeled auxin export assay (Lewis and Muday 2009; Tang et al. 2024). Fresh protonema tissues were used for all tested transformants. The result demonstrated that the auxin export activity of these hyperpolarized PpPINA variants was not impaired by mutations (Figure S2).

**Figure 5.**
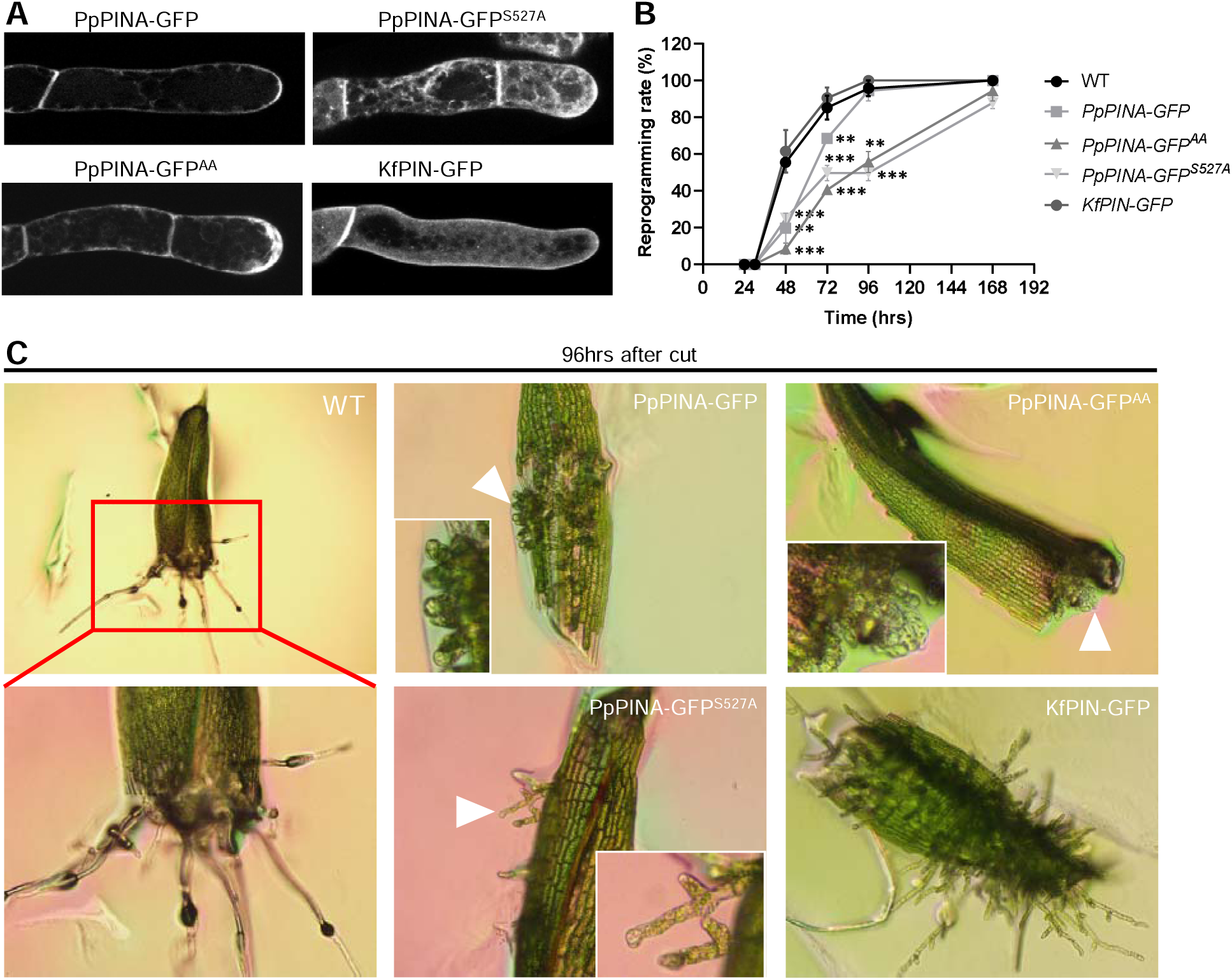
Overexpression of hyperpolarized PINs impedes moss leaf cell reprogramming. (A) Representative images of protonema tip cells in indicated overexpression background. In a wildtype background, the PIN-GFP constructs were inserted into a neutral locus driven by an inducible promoter. The tissues were cultured in an imaging dish for 5 days followed by 1 μM β-estradiol treatment for 3 days. The filaments were imaged by Leica confocal SP8. The signal of PpPINA mutants PpPINA-GFP^S527A^ and PpPINA-GFP^AA^ are hyperpolarized at the tip of protonema, causing swollen tips. The signal of KfPIN-GFP showed homogenized distribution on the plasma membrane. (B) The reprogramming rates of WT and indicated transgenic plants. Detached leaves from all examined plants were cultivated on the same solid media plate supplied with 1 μM β-estradiol. Overexpression of KfPIN-GFP did not affect cell reprogramming, while overexpression of PpPINA-GFP showed a slight delay in cell reprogramming. Overexpression of hyperpolarized PpPINA-GFP^S527A^ and PpPINA-GFP^AA^ revealed severe delayed cell reprogramming. (C) Example images of detached leaves from indicated transgenic plants. White arrowheads indicate tip cell protrusion. The mutants with overexpressed hyperpolarized PpPINA showed swollen tips, which confirmed the expressional induction after 4 days.

Next, we hypothesized that excessive hyperpolarized PpPINA variants would disrupt the polarity re-establishment during cell reprogramming. To test this, we examined the cell reprogramming rates of leaves detached from gametophores of the inducible overexpression lines carrying hyperpolarized PpPINA-GFP^S527A^ and PpPINA-GFP^AA^, respectively. Detached leaves were placed on a growth medium containing β-estradiol to constitutively induce gene expression of PpPINA variants. We found that hyperpolarized PpPINA-GFP variants accumulated at the site of future cell outgrowth, similar to WT PpPINA-GFP (Figure S3). Overexpression of hyperpolarized PpPINA-GFP variants significantly impaired reprogramming up to 96 hours (Figure 5B). Protonema cells regenerated from these hyperpolarized variants exhibited swollen tip phenotypes (Figure 5C, Figure S3), confirming successful overexpression.

To clarify that the defect is not caused by general overexpression of PpPINA, we generated the same inducible overexpression plants carrying WT forms of PpPINA-GFP and examined the reprogramming rate. The excess PpPINA-GFP revealed a slight delay of cell reprogramming at the first 72 hours, while most leaves were reprogrammed at 96 hours with similar efficiency as WT leaves.

To verify the importance of polarity, we included another non-polar KfPIN-GFP, obtained from the green algae *Klebsormidium flaccidum*, as an internal control. The KfPIN-GFP is known to export auxin but is evenly distributed across the plasma membrane in *P. patens* (Figure 5A, S2) (Skokan et al. 2019). Examination of reprogramming rates revealed that overexpression of non-polar KfPIN-GFP did not affect the reprogramming rate (Figure 5B, 5C). Additionally, we generated a phosphorylation mimic version of PpPINA^DD^-GFP under the same inducible control, which showed weak and even distribution on the plasma membrane and did not delay the cell reprogramming process (Figure S4). These results suggest that not only the proper amount but also polar, subcellular distribution of PpPINA are important for effective cell reprogramming.

## Discussion

### The contribution of auxin and PpPINA to moss leaf cell reprogramming

In this study, we discovered that auxin plays a minor role in cell reprogramming, contrary to its significant role in higher plant species where it promotes cell regeneration (Ishida et al. 2022; Mazur et al. 2020). Nonetheless, PpPINA, along with PpPINB, is crucial for this process. Time-lapse imaging showed that PpPINA accumulates in a polar manner at the site of future cell protrusion, suggesting it serves as a polarity landmark. This was further supported by experiments where hyperpolarized PpPINA variants delayed cell reprogramming, while WT PpPINA and non-polar KfPIN did not. These findings highlight PpPINA’s important role in establishing polarity in addition to auxin export, providing new insights into the mechanisms of cell reprogramming in *P. patens*.

In Arabidopsis, the polarization patterns of different PINs directly guide auxin flow which is crucial for almost all aspects of growth and development (Petrášek and Friml 2009). The significance of PIN for polarity regulation is tightly coupled with its function on auxin efflux (Adamowski and Friml 2015; Wiśniewska et al. 2006). In light of these observations, our findings lead to a re-evaluation of existing concepts considering PIN-related polarization. Based on the accumulation of PpPINA-GFP at polar sites preceding visible cell outgrowth, we propose a novel role of polarized PIN as a positional marker for cell reprogramming events.

### The auxin fluctuation is not completely correlated with the PpPINA accumulation

Given that PIN proteins are primarily known in all examined organisms from bryophytes to flowering plants for their role in auxin export, we investigated the relationship between auxin levels and PpPINA accumulation throughout cell reprogramming. In line with a previous study carried out in Arabidopsis, local auxin accumulation was shown in adjacent cells next to laser-ablated root cortex cells, around 3 hours before a restorative cell division (Hoermayer et al. 2020). However, the PIN localization in this process was not monitored.

In the context of regeneration of excised moss leaves, we observed that the amount of intracellular auxin did not necessarily mirror the expression levels of PpPINA (Figure 2, 4). During the cell reprogramming, auxin begins to accumulate approximately 24 hours post-cutting. An increase in subcellular auxin levels continues until the cell outgrowth becomes visible (Figure 2). In contrast, PpPINA exhibits specific polarization around 8 hours prior to visible outgrowth, which occurs approximately 36 hours post-cutting. In other words, regenerative cells continuously accumulate auxin around 8 hours before the cell outgrows when PpPINA expression and polarization become apparent (Figure S5). Our results revealed that overall auxin fluctuation is not completely correlated with the PpPINA accumulation, while only within 8 hours before cell outgrowing, both auxin levels and PpPINA expression increased thereby showing a temporary positive correlation. Although the primary function of PpPINA is to export auxin, the positive correlation between auxin levels and PpPINA expression and polarization suggests that the elevated auxin levels may result from increased local auxin biosynthesis.

In Arabidopsis, local auxin synthesis and PIN action work together to regulate subcellular auxin levels (Brumos et al. 2018; Petrášek and Friml 2009). Near the cutting sites of Arabidopsis root tips, rapid local auxin biosynthesis is triggered by multiple factors, initiating root regeneration but PIN-mediated auxin transport is not essential for these processes (Matosevich et al. 2020). This supports the idea that increased auxin biosynthesis might explain why the increase in auxin is not attenuated by the accumulation of PpPINA in moss reprogramming cells.

Additionally, we confirmed the PIN auxin export activity in filamentous protonema cells (Figure S2). Due to limited cell numbers, we could not validate the auxin export activity directly during cell reprogramming. Previous research from Arabidopsis indicates that PIN proteins need different kinases e.g. PINOID and D6 protein kinases to activate their export activity (Zourelidou et al. 2014). Thus, the temporary positive correlation between increased auxin levels and PpPINA accumulation at the later time frames of cell reprogramming suggests that polarized PpPINA may not be fully activated for auxin export. We reasoned that if PpPINA is expressed and fully activated, the auxin level is supposed to be decreased, but we see the opposite effect. This supports the hypothesis that the PpPINA in cell reprogramming is not solely linked to its auxin transport activity but may have additional roles.

### The proper polarity of PINA facilitates effective cell reprogramming

Regeneration patterns vary significantly among different organisms. In Arabidopsis, vascular tissue repair and root regeneration occur through new cell divisions and re-differentiation into new cell types (Hoermayer et al. 2020; Liu et al. 2014; Sauer et al. 2006). For instance, root epidermal cells adjacent to a local injury would trigger restorative cell division such that the newly divided cells regain the identity of the cortex and epidermal cells to restore the ablated cell (Hoermayer et al. 2020). In this process, newly formed cells typically do not undergo extensive cell expansion. In contrast, *P. patens* exhibits a distinct regeneration mechanism where leaf cells reprogram into filamentous stem cells through substantial cell expansion, ultimately forming new apical cells of a filament (Kofuji and Hasebe 2014).

In this context, intracellular polarity must be re-established to ensure that reprogramming cells grow in the correct direction. Our observations indicate that PpPINA polarizes well before visible cell outgrowth begins, suggesting that PpPINA plays a crucial role in re-establishing polarity and may act as a polarity marker. PpPINA may be recruited by other, not yet unidentified components, which guide its polarization to specific locations. Future research will focus on elucidating how PpPINA is recruited and on its biological function in the context of polarized cell outgrowth. Understanding these mechanisms will enhance our knowledge of polarity establishment in plant cell regeneration.

## Material and methods

### Plant growth and transformation

All transgenic plants used in this study are listed in Supplemental Table 1.

*P. patens* transgenic plants expressing pACT::PINA, pACT::PINB-GFP, PpPINA::PINA-GFP, and *pina, pinb*, and *pina pinb* were generated and verified as reported previously (Viaene et al. 2014). The *tara tarc* mutant was generated as published (Landberg et al. 2021). The auxin reporter line PpR2D2 was generated and the auxin responses were verified and reported in a previous study (Thelander et al. 2019). The inducible overexpression XVE::PpPINA-GFP, XVE:: PpPINA-GFP^S527A^, XVE::PpPINA-GFP^AA,^ and XVE::KfPIN-GFP lines were generated as described (Nishiyama et al. 2000; Yamada et al. 2016). Two independent transgenic lines were selected and verified for imaging and analysis. One representative line was shown in the results.

### Plasmid construction

The primers used in this study are listed in Supplemental Table 2. For transgenic moss lines with inducible overexpression, the PIN-GFP regions were amplified from previously generated plasmids, which used the genomic DNA for PpPINA and the coding sequence for KfPIN (Zhang et al. 2019). PIN-GFP was cloned into the Gateway entry plasmid pENTR/D-TOPO as the manufacturer suggested and subcloned into the pPGX8 vector, which contained a p35S-driven β-estradiol inducible XVE cassette via a Gateway LR reaction (Invitrogen) (Nakaoka et al. 2012).

### Microscopy analysis

Each transgenic line was observed every day until the seventh day post-cutting under a bright field of the BX53 biological microscope (Olympus). Most observations were performed with UIS2 Plan Achromat 20x/0.4 objective lens. The images were captured with the CellSens imaging software. In brief, after picking up gametophores, the top two young leaves and the bottom old leaves were removed with a razor blade. The remaining leaves were excised from the gametophores, with the cut made at approximately two-thirds of the leaf length from the tip. All harvested leaves were placed on a BCDAT growth medium for the following imaging.

For the time-lapse movies throughout cell reprogramming, the excised leaves containing PpR2D2 and proPINA::PpPINA-GFP were cultivated in one-well chamber (Thermo Scientific, Lab-tek Cat 155361) covered with a thin layer of BCDAT medium, and the chamber was observed with vertical-stage laser scanning confocal microscope (Zeiss 800), as described (Von Wangenheim et al. 2017). Live-cell imaging was taken over time up to 40 hours until the cell outgrow was visualized. Most images were taken with a Plan APOCHROMAT 20x/0.8 (420650-9901) objective; the DII-nTdTOMATO and autofluorescence of chloroplasts were excited by the 561 nM, MDII-nVENUS and GFP with the 488 nm laser line.

For the inducible overexpression lines, moss protonemata were cultured in glass-bottom dishes covered with BCD agar medium for 6-7 days before microscopy. 1μM β-estradiol diluted in liquid BCD medium was applied to the tissues for additional 3 days. Live-cell imaging was performed using a Leica SP8X-SMD confocal microscope equipped with a hybrid single-molecule detector (HyD) and an ultrashort pulsed white-light laser (50%; 1 ps at 40-MHz frequency). Leica Application Suite X was used for microscope control, and an HC PL APO CS2 403/1.20 water immersion objective was used for observing the samples. The time-gating system was activated to avoid autofluorescence emitted by chloroplasts.

### *P. patens* auxin export assay

The auxin export assay performed with transgenic moss plants was modified from the protocol developed for Arabidopsis seedlings (Lewis and Muday 2009). The protocol for moss was described in reference (Tang et al. 2024).

### Image quantification

The intensity of PpR2D2 signals with MDII-nVENUS and DII-nTdTOMATO was measured with the TrackMate function in the Fiji application. The function automatically detected and measured the signals over time. The gray values of each time point were recorded and the average intensity and ratio between MDII-nVENUS and DII-nTdTOMATO were calculated and plotted in Excel.

The accumulation of proPINA::PpPINA-GFP signals during cell reprogramming was measured in Fiji. Based on the accumulation, a small area was cropped and mean intensity was measured over time. The same crop area was moved to a non-regenerative site, where the mean intensity was quantified as a negative control.

### Measurement of reprogramming rate

The reprogramming rate was measured as described before (Ishikawa et al. 2019). Excised leaves were cultivated on a BCDAT growth medium and observed every 24 hours. Once cell outgrowing was visualized, it was recorded as a single event. The ratio of the number of leaves with at least one outgrowing event over the number of total leaves was quantified as the cell reprogramming rate at each time point.

## Data Availability

The data underlying this article are available in the article and in its online supplementary material.

## Acknowledgments

The authors sincerely thank Dr. Barbara Kloeckener Gruissem’s time and efforts in critical reading and constructive advice on the manuscript.

## Author contributions

H.T. and J.F. initiated and designed the experiments. H.T. carried out the quantitative analysis and wrote the manuscript. L.H.C. carried out the quantitative analysis.

## Reference

Adamowski, M. and Friml, J. (2015) PIN-dependent auxin transport: action, regulation, and evolution. The Plant Cell 27: 20–32.

Birnbaum, K.D. and Alvarado, A.S. (2008) Slicing across kingdoms: regeneration in plants and animals. Cell 132: 697–710.

Brumos, J., Robles, L.M., Yun, J., Vu, T.C., Jackson, S., Alonso, J.M., et al. (2018) Local auxin biosynthesis is a key regulator of plant development. Developmental cell 47: 306–318. e305.

Gallei, M., Luschnig, C. and Friml, J. (2020) Auxin signalling in growth: Schrödinger’s cat out of the bag. Current opinion in plant biology 53: 43–49.

Hajný, J., Tan, S. and Friml, J. (2022) Auxin canalization: From speculative models toward molecular players. Current opinion in plant biology 65: 102174.

Hoermayer, L., Montesinos, J.C., Marhava, P., Benková, E., Yoshida, S. and Friml, J. (2020) Wounding-induced changes in cellular pressure and localized auxin signalling spatially coordinate restorative divisions in roots. Proceedings of the National Academy of Sciences 117: 15322–15331.

Ikeuchi, M., Favero, D.S., Sakamoto, Y., Iwase, A., Coleman, D., Rymen, B., et al. (2019) Molecular mechanisms of plant regeneration. Annual review of plant biology 70: 377–406.

Ikeuchi, M., Ogawa, Y., Iwase, A. and Sugimoto, K. (2016) Plant regeneration: cellular origins and molecular mechanisms. Development 143: 1442–1451.

Ishida, S., Suzuki, H., Iwaki, A., Kawamura, S., Yamaoka, S., Kojima, M., et al. (2022) Diminished auxin signaling triggers cellular reprogramming by inducing a regeneration factor in the liverwort Marchantia polymorpha. Plant and Cell Physiology 63: 384–400.

Ishikawa, M., Morishita, M., Higuchi, Y., Ichikawa, S., Ishikawa, T., Nishiyama, T., et al. (2019) Physcomitrella STEMIN transcription factor induces stem cell formation with epigenetic reprogramming. Nature Plants 5: 681–690.

Ishikawa, M., Murata, T., Sato, Y., Nishiyama, T., Hiwatashi, Y., Imai, A., et al. (2011) Physcomitrella cyclin-dependent kinase A links cell cycle reactivation to other cellular changes during reprogramming of leaf cells. The Plant Cell 23: 2924–2938.

Kofuji, R. and Hasebe, M. (2014) Eight types of stem cells in the life cycle of the moss Physcomitrella patens. Current opinion in plant biology 17: 13–21.

Landberg, K., Šimura, J., Ljung, K., Sundberg, E. and Thelander, M. (2021) Studies of moss reproductive development indicate that auxin biosynthesis in apical stem cells may constitute an ancestral function for focal growth control. New Phytologist 229: 845–860.

Lewis, D.R. and Muday, G.K. (2009) Measurement of auxin transport in Arabidopsis thaliana. Nature protocols 4: 437–451.

Li, C., Sako, Y., Imai, A., Nishiyama, T., Thompson, K., Kubo, M., et al. (2017) A Lin28 homologue reprograms differentiated cells to stem cells in the moss Physcomitrella patens. Nature communications 8: 14242.

Liu, J., Sheng, L., Xu, Y., Li, J., Yang, Z., Huang, H., et al. (2014) WOX11 and 12 are involved in the first-step cell fate transition during de novo root organogenesis in Arabidopsis. The Plant Cell 26: 1081–1093.

Marhava, P., Hoermayer, L., Yoshida, S., Marhavý, P., Benková, E. and Friml, J. (2019) Re-activation of stem cell pathways for pattern restoration in plant wound healing. Cell 177: 957–969. e913.

Matosevich, R., Cohen, I., Gil-Yarom, N., Modrego, A., Friedlander-Shani, L., Verna, C., et al. (2020) Local auxin biosynthesis is required for root regeneration after wounding. Nature Plants 6: 1020–1030.

Mazur, E., Kulik, I., Hajný, J. and Friml, J. (2020) Auxin canalization and vascular tissue formation by TIR1/AFB-mediated auxin signaling in Arabidopsis. New Phytologist 226: 1375–1383.

Nakaoka, Y., Miki, T., Fujioka, R., Uehara, R., Tomioka, A., Obuse, C., et al. (2012) An Inducible RNA Interference System in Physcomitrella patens Reveals a Dominant Role of Augmin in Phragmoplast Microtubule Generation The Plant Cell 24: 1478–1493.

Nishiyama, T., Hiwatashi, Y., Sakakibara, K., Kato, M. and Hasebe, M. (2000) Tagged Mutagenesis and Gene-trap in the Moss, Physcomitrella patens by Shuttle Mutagenesis. DNA Research 7: 9–17.

Petrášek, J. and Friml, J. (2009) Auxin transport routes in plant development.

Rensing, S.A., Goffinet, B., Meyberg, R., Wu, S.-Z. and Bezanilla, M. (2020) The Moss Physcomitrium (Physcomitrella) patens: a model organism for non-seed plants. The Plant Cell 32: 1361–1376.

Sauer, M., Balla, J., Luschnig, C., Wiśniewska, J., Reinöhl, V., Friml, J., et al. (2006) Canalization of auxin flow by Aux/IAA-ARF-dependent feedback regulation of PIN polarity. Genes & development 20: 2902–2911.

Skokan, R., Medvecká, E., Viaene, T., Vosolsobě, S., Zwiewka, M., Müller, K., et al. (2019) PIN-driven auxin transport emerged early in streptophyte evolution. Nature Plants 5: 1114–1119.

Sugimoto, K., Temman, H., Kadokura, S. and Matsunaga, S. (2019) To regenerate or not to regenerate: factors that drive plant regeneration. Current opinion in plant biology 47: 138–150.

Tang, H., Lu, K.-J., Zhang, Y., Cheng, Y.-L., Tu, S.-L. and Friml, J. (2024) Divergence of trafficking and polarization mechanisms for PIN auxin transporters during land plant evolution. Plant Communications 5.

Thelander, M., Landberg, K. and Sundberg, E. (2019) Minimal auxin sensing levels in vegetative moss stem cells revealed by a ratiometric reporter. New Phytologist 224: 775–788.

Viaene, T., Landberg, K., Thelander, M., Medvecka, E., Pederson, E., Feraru, E., et al. (2014) Directional auxin transport mechanisms in early diverging land plants. Current Biology 24: 2786–2791.

Von Wangenheim, D., Hauschild, R., Fendrych, M., Barone, V., Benková, E. and Friml, J. (2017) Live tracking of moving samples in confocal microscopy for vertically grown roots. Elife 6: e26792.

Wiśniewska, J., Xu, J., Seifertová, D., Brewer, P.B., Růžička, K., Blilou, I., et al. (2006) Polar PIN Localization Directs Auxin Flow in Plants. Science 312: 883–883.

Xu, L. (2018) De novo root regeneration from leaf explants: wounding, auxin, and cell fate transition. Current opinion in plant biology 41: 39–45.

Yamada, M., Miki, T. and Goshima, G. (2016) Imaging mitosis in the moss Physcomitrella patens. The Mitotic Spindle: Methods and Protocols: 263–282.

Ye, B.-B., Shang, G.-D., Pan, Y., Xu, Z.-G., Zhou, C.-M., Mao, Y.-B., et al. (2020) AP2/ERF transcription factors integrate age and wound signals for root regeneration. The Plant Cell 32: 226–241.

Zhang, Y., Xiao, G., Wang, X., Zhang, X. and Friml, J. (2019) Evolution of fast root gravitropism in seed plants. Nature communications 10: 3480.

Zourelidou, M., Absmanner, B., Weller, B., Barbosa, I.C., Willige, B.C., Fastner, A., et al. (2014) Auxin efflux by PIN-FORMED proteins is activated by two different protein kinases, D6 PROTEIN KINASE and PINOID. elife 3: e02860.

