## Supplemental Figures for "Auxin fluctuation and PIN polarization in moss leaf cell reprogramming"

### Slide 1
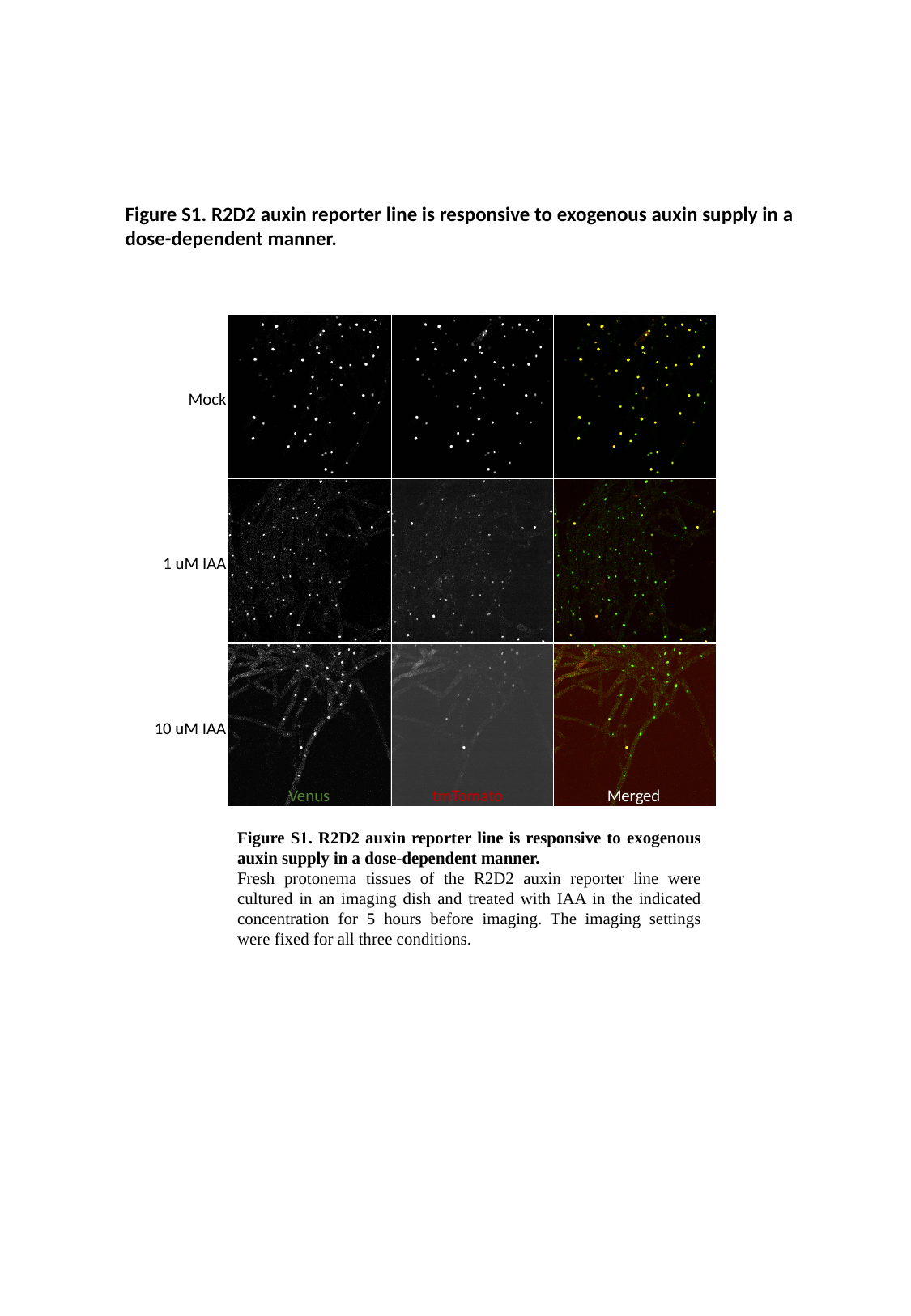

Figure S1. R2D2 auxin reporter line is responsive to exogenous auxin supply in a dose-dependent manner.
Mock
1 uM IAA
10 uM IAA
Venus
tmTomato
Merged
Figure S1. R2D2 auxin reporter line is responsive to exogenous auxin supply in a dose-dependent manner.
Fresh protonema tissues of the R2D2 auxin reporter line were cultured in an imaging dish and treated with IAA in the indicated concentration for 5 hours before imaging. The imaging settings were fixed for all three conditions.

### Slide 2
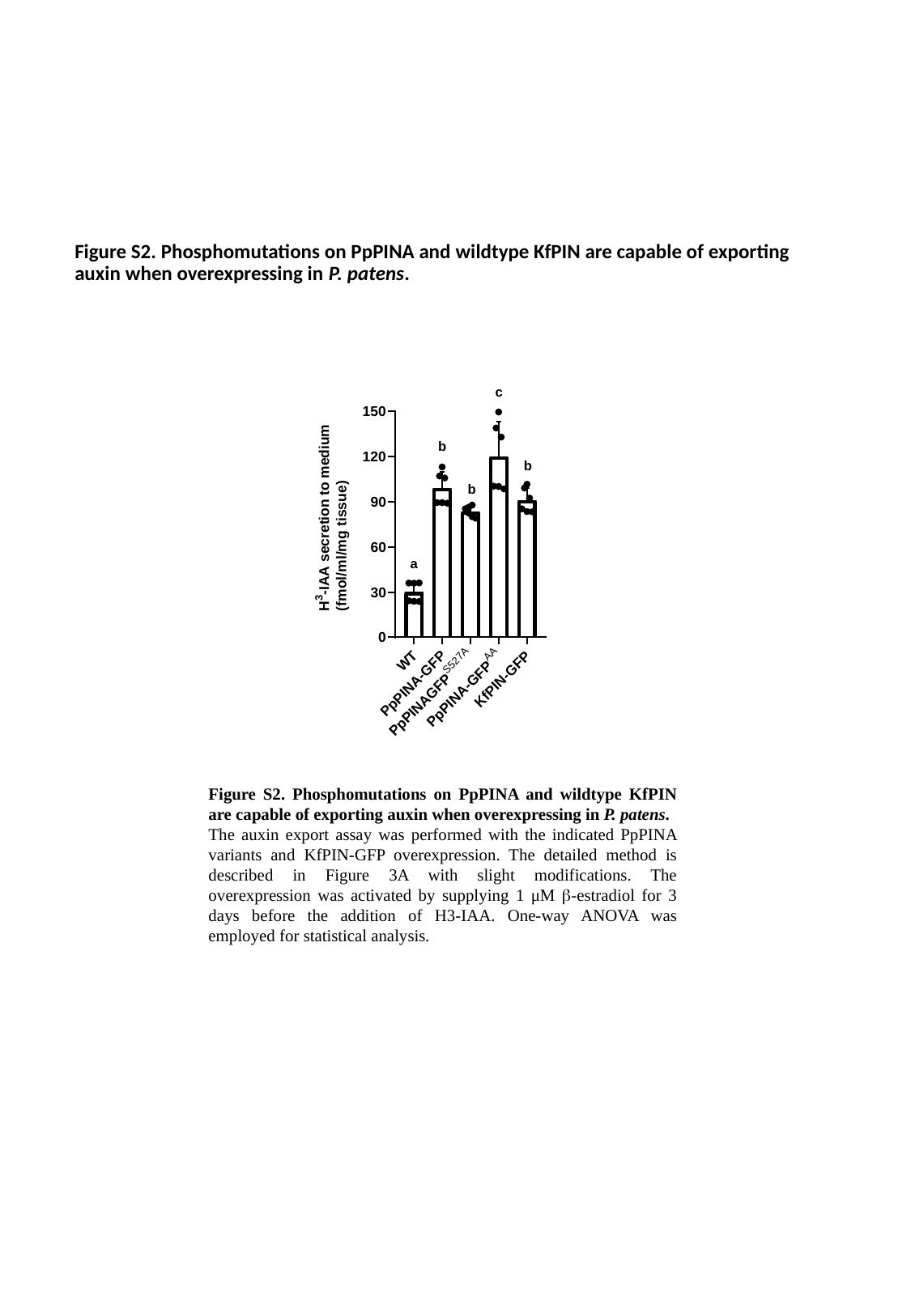

#### Figure S2. Phosphomutations on PpPINA and wildtype KfPIN are capable of exporting auxin when overexpressing in P. patens.
c
b
b
b
a
Figure S2. Phosphomutations on PpPINA and wildtype KfPIN are capable of exporting auxin when overexpressing in P. patens.
The auxin export assay was performed with the indicated PpPINA variants and KfPIN-GFP overexpression. The detailed method is described in Figure 3A with slight modifications. The overexpression was activated by supplying 1 μM β-estradiol for 3 days before the addition of H3-IAA. One-way ANOVA was employed for statistical analysis.

### Slide 3
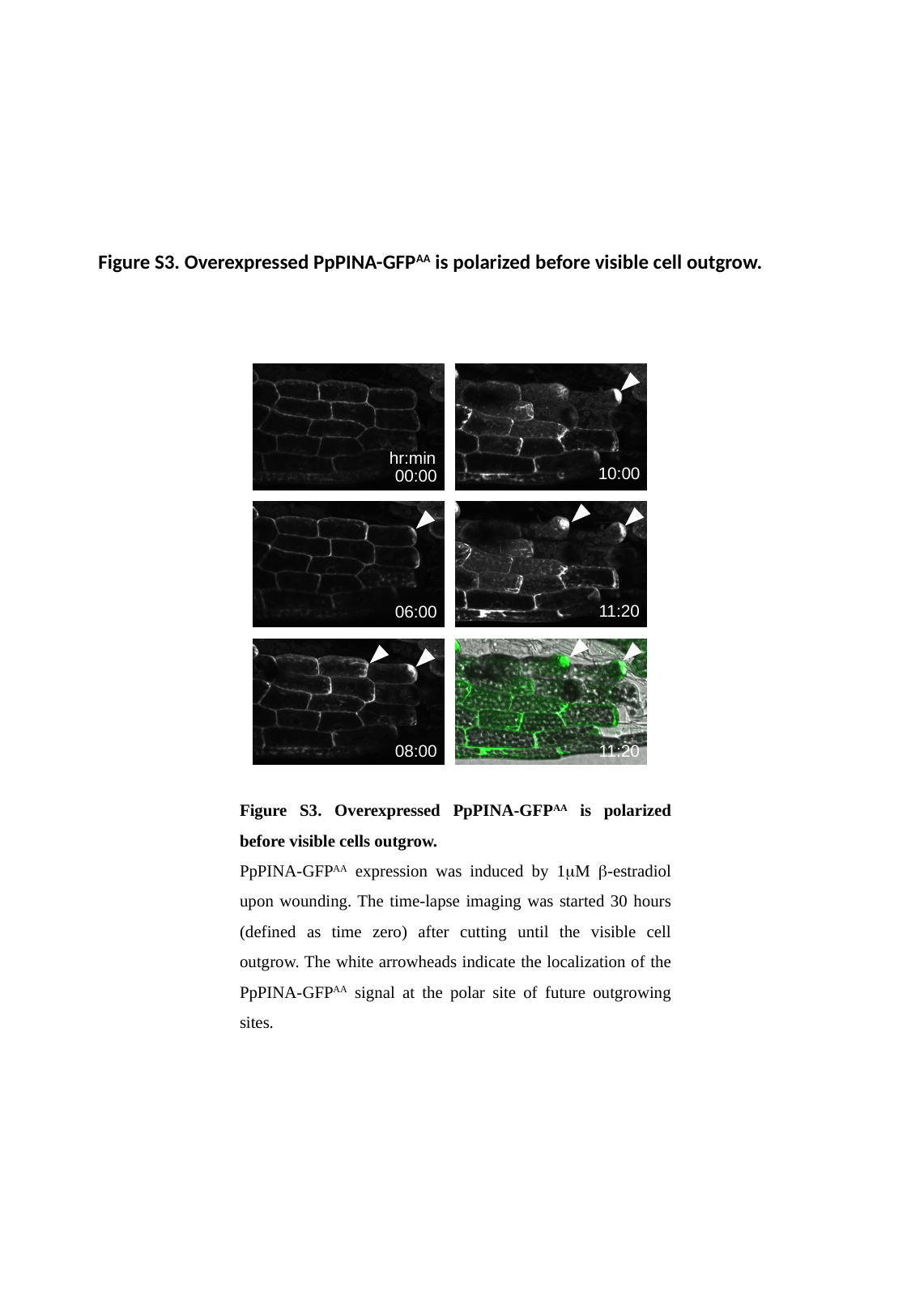

#### Figure S3. Overexpressed PpPINA-GFP^AA^ is polarized before visible cell outgrow.
hr:min
10:00
00:00
11:20
06:00
08:00
11:20
Figure S3. Overexpressed PpPINA-GFP^AA^ is polarized before visible cells outgrow.
PpPINA-GFP^AA^ expression was induced by 1μM β-estradiol upon wounding. The time-lapse imaging was started 30 hours (defined as time zero) after cutting until the visible cell outgrow. The white arrowheads indicate the localization of the PpPINA-GFP^AA^ signal at the polar site of future outgrowing sites.

### Slide 4
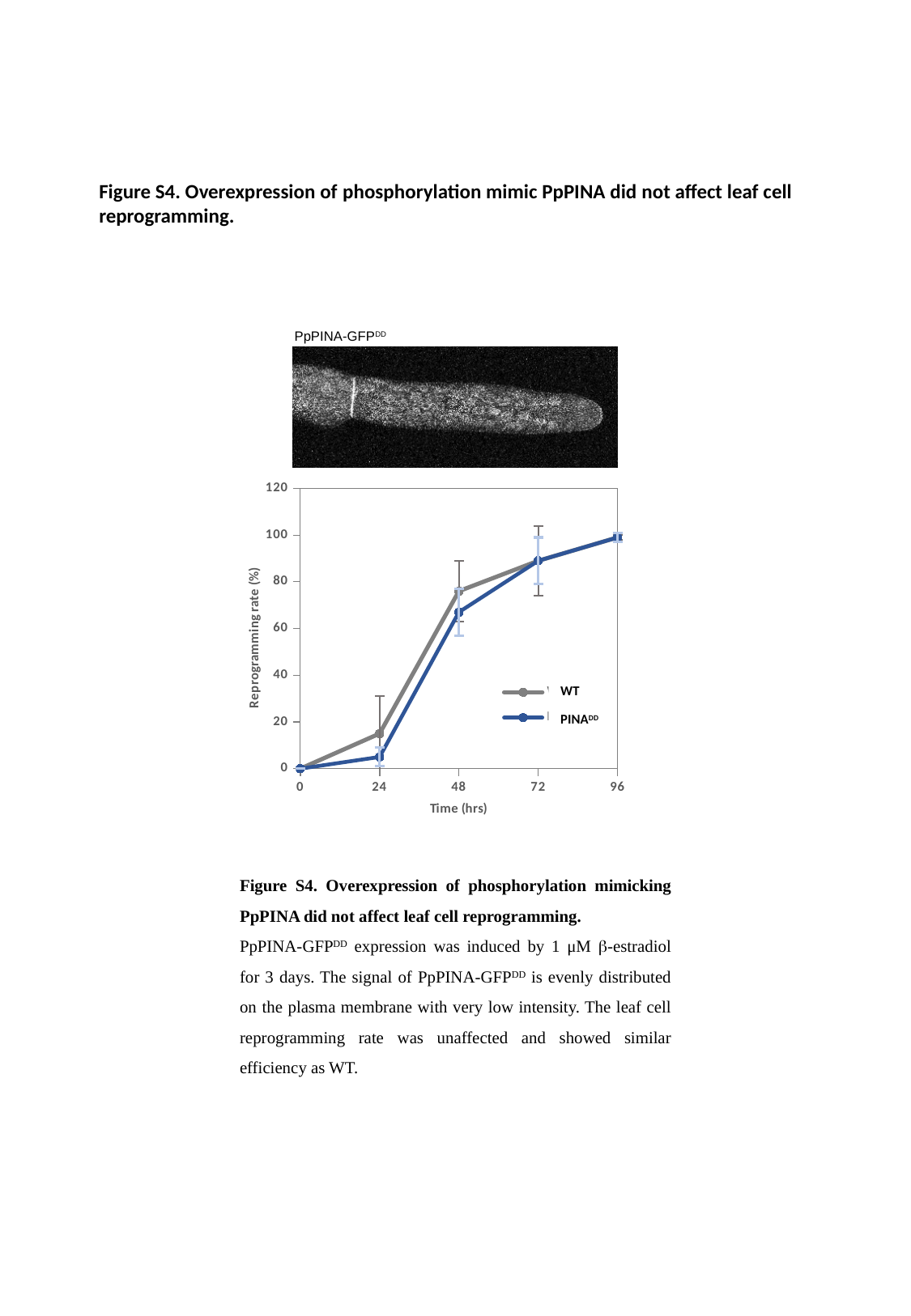

Figure S4. Overexpression of phosphorylation mimic PpPINA did not affect leaf cell reprogramming.
PpPINA-GFP^DD^
###### Chart
| Category | WT | PINADD |
|---|---|---|
| 0 | 0.0 | 0.0 |
| 24 | 15.0 | 5.0 |
| 48 | 76.0 | 67.0 |
| 72 | 89.0 | 89.0 |
| 96 | 99.0 | 99.0 |WT
PINA^DD^
Figure S4. Overexpression of phosphorylation mimicking PpPINA did not affect leaf cell reprogramming.
PpPINA-GFP^DD^ expression was induced by 1 μM β-estradiol for 3 days. The signal of PpPINA-GFP^DD^ is evenly distributed on the plasma membrane with very low intensity. The leaf cell reprogramming rate was unaffected and showed similar efficiency as WT.

### Slide 5
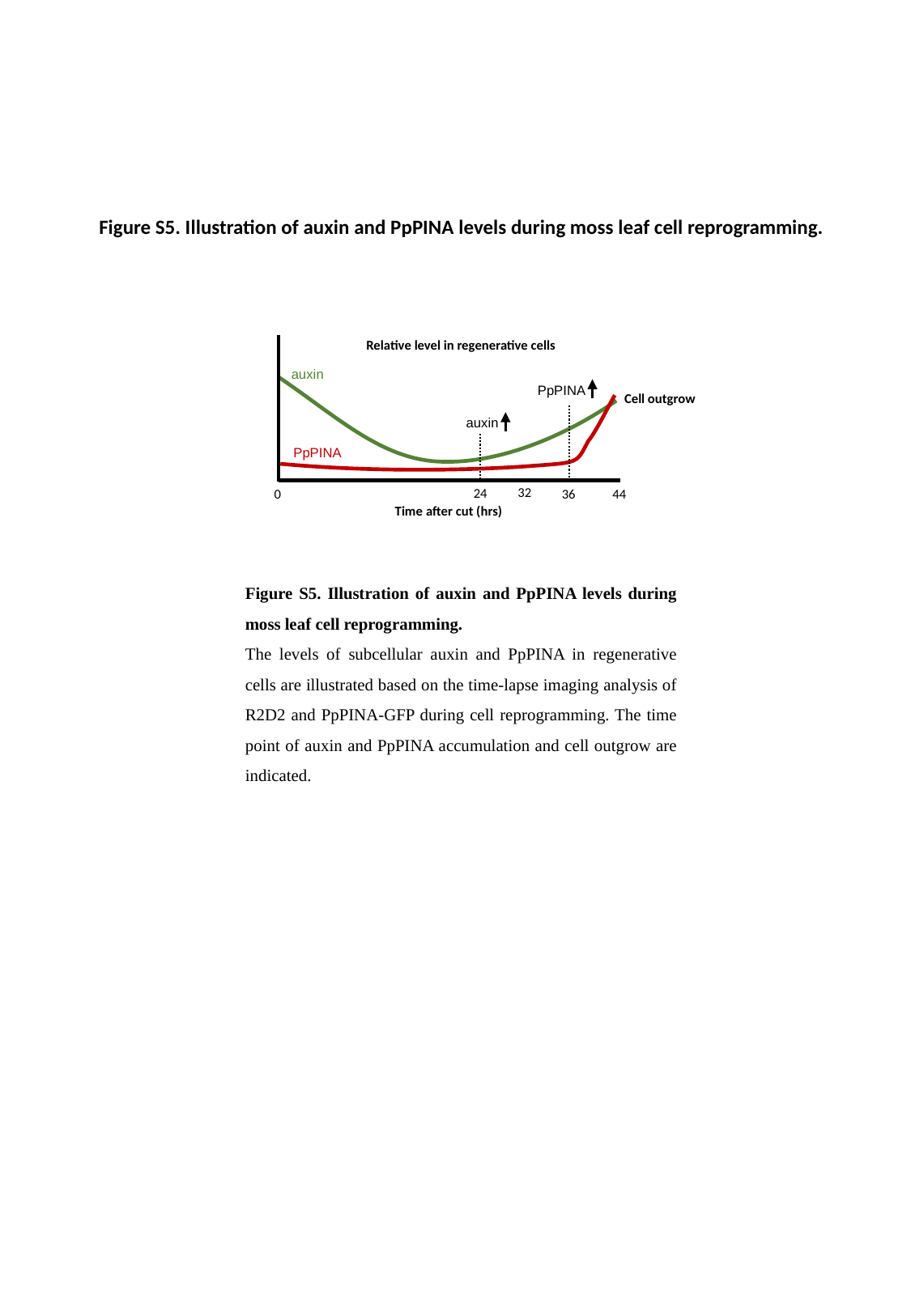

#
Figure S5. Illustration of auxin and PpPINA levels during moss leaf cell reprogramming.
Relative level in regenerative cells
auxin
PpPINA
Cell outgrow
auxin
PpPINA
32
24
44
0
36
Time after cut (hrs)
Figure S5. Illustration of auxin and PpPINA levels during moss leaf cell reprogramming.
The levels of subcellular auxin and PpPINA in regenerative cells are illustrated based on the time-lapse imaging analysis of R2D2 and PpPINA-GFP during cell reprogramming. The time point of auxin and PpPINA accumulation and cell outgrow are indicated.
